# The BacA(SbmA) Importer of Symbiotically Important Legume Nodule Cysteine-Rich Peptides: Insights into Protein Architecture, Function, and Evolutionary Implications

**DOI:** 10.1101/2025.09.17.676847

**Authors:** Markus F.F. Arnold, Siva Sankari, Michael Deutsch, Charley C. Gruber, Francisco J. Guerra-Garcia, Kostantinos Beis, Graham C. Walker

## Abstract

Some legumes encode families of NCR (Nodule-Cysteine-Rich) peptides that cause their rhizobial partners to terminally differentiate during the development of a nitrogen-fixing symbiosis. *Sinorhizobium meliloti*, whose plant hosts *Medicago truncatula* and *M. sativa* express *ca*. 600 NCR peptides during root nodule development, possesses a symbiotically essential BacA_*Sm*_ protein that imports certain NCR peptides into the cytoplasm. This import permits proteolytic degradation of the NCR peptides, thereby protecting the endocytosed bacteria from their antimicrobial peptide-like lethality, while also allowing certain NCR peptides to undergo their symbiotically critical interactions with cytoplasmic components, for example heme-sequestration in the case of NCR247. BacA’s *Escherichia coli* ortholog SbmA_*Ec*_ can restore a wildtype phenotype to a *ΔbacA*_*Sm*_ mutant. Our study employed 54 *S. meliloti bacA*_Sm_ missense mutants (35 to cysteine and 19 to glycine) that we tested for protein production, ability to establish a nitrogen-fixing symbiosis, and their susceptibility to killing by higher levels of the NCR247 and the Bac7(1-35) peptides. We also used the Single Cysteine Accessibility Method to make topological inferences. Our detailed genetic, biochemical, structural, and physiological analyses have revealed that BacA_Sm_ and SbmA homodimers function as finely tuned import machines, whose structures can be relatively easily disrupted by single amino acid changes. Our discovery that several mutations that differentially separate nitrogen-fixation, NCR247 import, and Bac7(1-35) import map to the lining of the peptide-binding cavity in the outward-open SbmA/BacA conformation suggests a molecular explanation the other otherwise paradoxical observation that SbmA/BacAs from pathogens can fully replace BacA_*Sm*_, whereas BacAs from other rhizobia cannot.

**Significance Statement:** *Sinorhizobium meliloti* BacA_*Sm*_ and *Escherichia coli* SbmA_*Ec*_ are closely related proteins that function as homodimeric transporters to import peptides and other cargos through the cytoplasmic membrane into the cytoplasm. BacA is critical for *S. meliloti* to establish a nitrogen-fixing symbiosis with its legume hosts because of its ability to import Nodule Cysteine-Rich (NCR) nodule-specific plant peptides. This import protects the bacteria inside the nodule from the potentially lethal effects of these NCR peptides while also enabling NCRs to make their intracellular interactions that are necessary for symbiosis. Our extensive multidisciplinary studies offer new insights into function of BacA/SbmA transporters and provide a molecular explanation for why BacA/SbmA orthologs from mammalian pathogens can replace BacA_*Sm*_ but those from other rhizobia cannot.

## Introduction

Legumes can grow without nitrogen fertilizer because they are able to establish a symbiosis with rhizobia in which the bacteria convert N_2_ gas into ammonia (nitrogen-fixation) in return for fixed carbon from the plant. After invading the root nodules they elicit, the rhizobia are endocytosed into membrane compartments (symbiosomes) within the cytoplasm of plant cells in the interior of the nodule where they undergo a set of physiological changes that convert the rhizobia into nitrogen-fixing bacteroids (1). In some legumes, the plant utilizes a family of defensin-like Nodule Cysteine Rich (NCR) peptides expressed specifically in the nodules to cause the endocytosed bacteria to terminally differentiate, a process thought to increase the efficiency of nitrogen fixation (2-4). In the case of the *Sinorhizobium meliloti*/*Medicago truncatula* symbiosis, the plant NCR peptide family consists of more than 600 cationic, acidic, and neutral members ranging from 24-65 amino acids, which are expressed in successive waves as the symbiosis is being established (2, 5-9). A few NCR peptides have shown to be critically required to establish the symbiosis (10-13), whereas others are not (14), and still others play a role determining fixation level compatibility between the host and the symbiont (15, 16). The detection of *ca*. 140 NCR peptides in the cytosol of bacteroids is consistent with some of the NCR peptides also interacting with intracellular bacterial targets (17, 18). At higher concentrations, cationic NCR peptides with pI >9 can act as fast-acting antibiotics that can kill bacteria through disruption of their cytoplasmic membrane (19, 20).

The smallest and best characterized NCR peptide is the 24 amino-acid NCR247 from *M. truncatula*, which is essential for symbiosis (13). NCR247 binds heme with nanomolar affinity and sequesters it, first into hexamers and then higher order complexes that render the heme biologically unavailable (13). Once imported into the cytoplasm of developing bacteroids, NCR247 creates a physiological state of heme deprivation, which in turn induces the import of the high levels of iron necessary for nitrogenase synthesis (13, 21, 22). Prior observations have shown that NCR247 also interacts with multiple proteins (21, 22) We have recently shown that, besides acting in the cytoplasm to induce iron import, NCR247 acts in the periplasm to induce the ChvI/ExoS- and FeuP/FeuQ-controlled regulons (Sankari *et al*., BioRxiv 2025).

The rhizobial BacA protein (and in some cases a BacA-like protein) plays essential roles in symbioses with legumes that express NCR peptides. *S. meliloti bacA* (BacA_*Sm*_) null mutants elicit ineffective nodules containing bacteria within the infection threads but no mature bacteroids because the rhizobia are killed upon release from the infection thread (23, 24). BacA is an inner membrane protein that is strongly homologous to the *Escherichia coli* SbmA (SbmA_*Ec*_) protein (24), which has been principally studied for its role in importing structurally unrelated peptides such as microcin B17, peptide derivatives, and aminoglycosides (25). Despite *E. coli* not being a plant symbiont, SbmA_*Ec*_ proved to be isofunctional with *S. meliloti* BacA_*Sm*_ (26), completely suppressing the symbiotic deficiencies of an *S. meliloti bacA* mutant, as did the BacA_*Ba*_ ortholog of the intracellular mammalian pathogen *Brucella abortus* (27). The *bacA*^*+*^ genes from a variety of rhizobia failed to restore nitrogen fixation to an *S. meliloti ΔbacA* mutant (28-31), although they did allow some limited developmental progression towards the bacteroid state.

Gail Ferguson and her colleagues elucidated a major symbiotic role for BacA when they showed that it plays a critical role in protecting *S. meliloti* from the antimicrobial effects of the *M. truncatula* NCR peptides (32, 33). Both the BacA_*Sm*_, the orthologous SbmA_*Ec*_, and BacA_*Ba*_ make the bacteria resistant to killing by higher levels of the NCR247 peptide and Bac7 (1-35) peptides by importing them into the cytoplasm where they can be degraded (34, 35). In addition, BacA-mediated import of certain NCR peptides allows them to interact with their cytoplasmic targets, for example heme in the case of NCR247 (13, 22).

The cryo-EM structures of both the SbmA_*Ec*_ and BacA_*Sm*_ transporters in the outward-open conformation revealed a homodimeric structure with a novel fold termed the SbmA-Like Peptide Transporter fold (SLiPT) (25, 36). The structure consists of a core transmembrane domain (TMD), comprised of 12 TMs (six from each protomer), while two additional TM0 domains (formed of TM0a and TM0b helices) flank the TMD, forming two peripheral domains (25). Although the overall TMD structure resembles that of type IV ABC transporters, SbmA_*Ec*_ and BacA_*Sm*_ are instead energized by the proton gradient with the proton translocation pathway defined by a glutamate ladder that is formed by conserved glutamates from both protomers within the TMD (25, 36); the structures revealed a central gate that isolates the peptide-binding cavity from the glutamate ladder in the outward-open conformation. The structure of both the SbmA_*Ec*_ and BacA_*Sm*_ was determined in an outward-open conformation, which has the cavity open to the periplasm (25). Subsequently, we solved the structure of SbmA_*Ec*_ in the inward-open conformation that has the cavity facing the cytoplasm (36). Opening of the central gate (Y372 in SbmA_*Ec*_; Y368 in BacA_*Sm*_) and movement of the proton along the glutamate ladder changes the shape of the cavity from an hourglass in the outward-open conformation to a cone shape in the inward-open conformation.

In this work we used genetic, biochemical, physiological, and structural analysis to gain significant insights into how BacA’s structure and conformational changes enable it to carry out its critical complex roles in the *Rhizobium*-legume symbiosis. In addition, our observations have allowed us to propose a model to explain how rhizobial BacAs can import a wide variety of NCR peptides yet undergo evolutionary tuning to match the specific family of NCRs made by their legume host. This model also suggests an explanation for the paradox that BacA orthologs from the mammalian pathogens *B. abortus and E. coli* are isofunctional with *S. meliloti* BacA,yet BacA orthologs from various other rhizobia are not.

## Results

### Summary of the study

Our study employed 54 *S. meliloti bacA*_Sm_ missense mutants (Fig. 1): 35 newly constructed single cysteine *bacA*_Sm_ mutants and 19 single glycine *bacA*_Sm_ mutants that had been partially characterized in an earlier publication prior to our awareness of NCR peptides (37). Since BacA_Sm_ lacks cysteines despite being 420 amino acids in length, we chose to make monocysteine mutants so we could employ the Single Cysteine Accessibility Method (SCAM) to make inferences about the topology of the BacA protein in living cells under different conditions (38). As summarized in Fig.1, we used *S. meliloti ΔbacA* derivatives carrying plasmids with the various *bacA* alleles to test: i) whether the mutants made BacA_Sm_ protein or not, ii) their ability to establish a nitrogen-fixing symbiosis (Table S4), iii) their susceptibility to killing by higher levels of the NCR247 peptide (Fig. S1), and iv) their susceptibility to killing by the Bac7 (1-35) peptide (Fig. S2). Examples of unambiguous topological information inferred from SCAM analyses of the single cysteine *bacA* mutants that made BacA_Sm_ protein (Fig. S3) are shown in (Fig. 1 middle left). Our inferences about the topology of BacA *in vivo* based on our SCAM analyses were all in complete agreement with the published BacA and SbmA structures.

**Figure 1.**
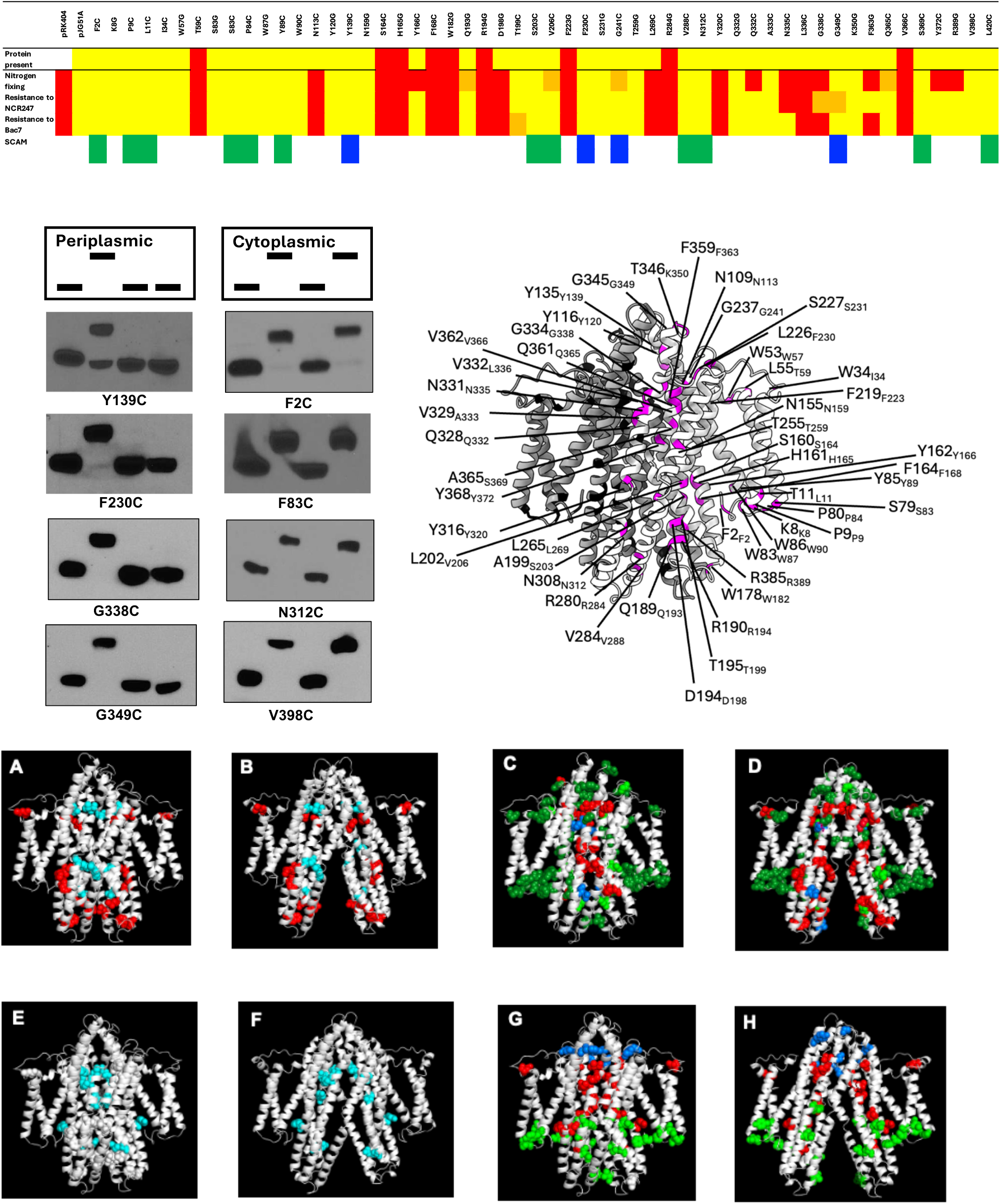
Properties and phenotypes of the 54 Gly and Cys missense *bacA*_Sm_ mutants of *S. meliloti* displayed at the corresponding positions of the open-outward and open-inward conformations of its homologous isofunctional *E*. coli ortholog, SbmA_Ec_. The Summary at the top shows the properties and phenotypes properties of the BacA_Sm_ missense mutants with respect to: i) whether they made BacA_Sm_ protein detectable in Western blots protein or not, ii) their ability to establish a nitrogen-fixing symbiosis, iii) their susceptibility to killing by higher levels of the NCR247 peptide, iv) their susceptibility to killing by higher levels of the Bac7(1-35) peptide, and v) the topological location of certain BacA monocysteine mutants with a wild type phenotypes inferred by SCAM methodology (examples shown middle left). The middle right shows the SbmA_*Ec*_ equivalents of the 54 BacA_*Sm*_ mutants we studied displayed one of the subunits of the SbmA_*Ec*_ homodimeric transporter. The *E. coli* SbmA numbering is shown with the corresponding *S. meliloti* BacA amino acid numbering subscripted. (**A, B**). 14 of the 54 *bacA*_Sm_ mutants exhibited a null (Fix^-^ NCR247^-^ Bac7^-^) or virtually null phenotype. The locations of these 14 altered amino acids in the mutant BacA_Sm_ proteins are shown at the corresponding locations on the outward-open (**A**) and inward-open SbmA_Ec_ structures (**B**). The positions of the 9 mutated amino acids that result in no protein detectable in Western blots are shown in red. The positions of the 5 mutated amino acids that do make protein are shown in cyan. (**C**,**D**). The locations of the 45 missense BacA mutants that do make protein with respect to their Fix^+^ phenotype (Fix^+^ dark green, Fix^+/-^ light green, Fix^-^ red, mixed blue) are shown at the corresponding locations on the outward-open (**C**) and inward-open (**D**) SbmA_Ec_ structures. **(E**,**F**) The location of *bacA* mutants that make protein but exhibit a partial-loss-of-function phenotype (“split phenotype”) are shown at the corresponding locations on the outward-open (**E**) and inward-open (**F)** SbmA_Ec_ structures. (**G, H**). The location of the 31 missense BacA_Sm_ Cys mutants that make protein are shown at the corresponding locations on the outward-open **(G**) and inward-open (**H**) SbmA_Ec_ structures. The color indicates the results of efforts to determine their *in vivo* topological position using the SCAM procedure: cytoplasmic (green), periplasmic (blue), anomalous (non-canonical) SCAM result (red).

As summarized in (Fig. 1 A-H), we are now able to interpret the phenotypic effects of the amino acid changes in these 54 missense BacA_*Sm*_ proteins in terms of their structure. Since the resolution of the BacA_*Sm*_ structure is limited to 6 Å, we have chosen to display the positions of the 54 BacA changes at the corresponding positions of the more highly resolved SbmA_*Ec*_ outward-open (Fig. 1 middle right) shows these correspondences) and inward-open structures (25). Because SbmA_*Ec*_ and BacA_*Sm*_ are so strongly closely related (Fig. S4) that they are isofunctional (26), most of our structural inferences are likely to be correct but we note this limitation. In principle, the amino acid changes we introduced could perturb BacA_*Sm*_ membrane insertion, folding, dimerization, binding to the peptide cargo, or the conformational changes necessary for peptide transport, which involve the movement of numerous helices and the making and breaking of multiple amino acids. Collectively, our analyses indicate that BacA_*Sm*_ and SbmA_*Ec*_ function as finely tuned import machines, whose structures can be relatively easily disrupted by single amino acid changes to cysteines or glycines in ways that either prevent a stable protein from being made or else result in an intact but non-functional protein. Other single amino acid mutations had more subtle effects on BacA function that proved to be informative. We corroborate the phenotypic behavior of a few distinct mutants in light of our published cryo-EM structures (25, 36).

### Missense mutations that result in a null phenotype and no protein

Fourteen of the 54 missense mutants had a null (Fix^-^ NCR247^-^ Bac7^-^) or virtually null phenotype. Of these, nine make no or virtually no protein (BacA_*Sm*_ mutant/SbmA_*Ec*_: T59C/L55, S164C/S160, H165G/H161, F168C/F164, W182G/W178, R194G/R190, F223G/F219, R284G/R280, V366C/362) presumably because the amino acid substitution makes the mutant BacA susceptible to proteolytic degradation by preventing proper membrane insertion, proper folding, or dimerization. Consideration of the location of these amino acid changes in the outward facing and inward facing structures offers insights into why the mutant proteins may result in unstable or partially folded proteins that would be susceptible to proteolysis. For example, as shown in Fig. 2A, the T59C/L55 and F223G/F219 mutations are within the TM0 domain PG lipid interface vicinity that are likely to interfere with the correct insertion to the membrane; TM0 does not undergo conformational changes during the transport cycle and it has been suggested to anchor the TMD to the inner membrane, therefore small changes in sequence are not tolerated (25, 36). As shown in Fig. 2B, F168/F164, which is located in the TM2 helix of one subunit, interacts with an aromatic rich pocket formed by Y313’/Y309’, F314’/F310’, and Y309’/Y313’ of the TM5’ in the other protomer. This cluster of amino acids is present in both the outward- and inward-open conformations and likely helps to stabilize the dimer interface during the various conformational changes associated with transport. It seems likely mutation of the nearby residues on the TM2 helix, S164/S160, H165/H161 to cysteine and glycine, respectively, similarly interferes with stable dimer formation (Fig. 2B).

**Figure 2.**
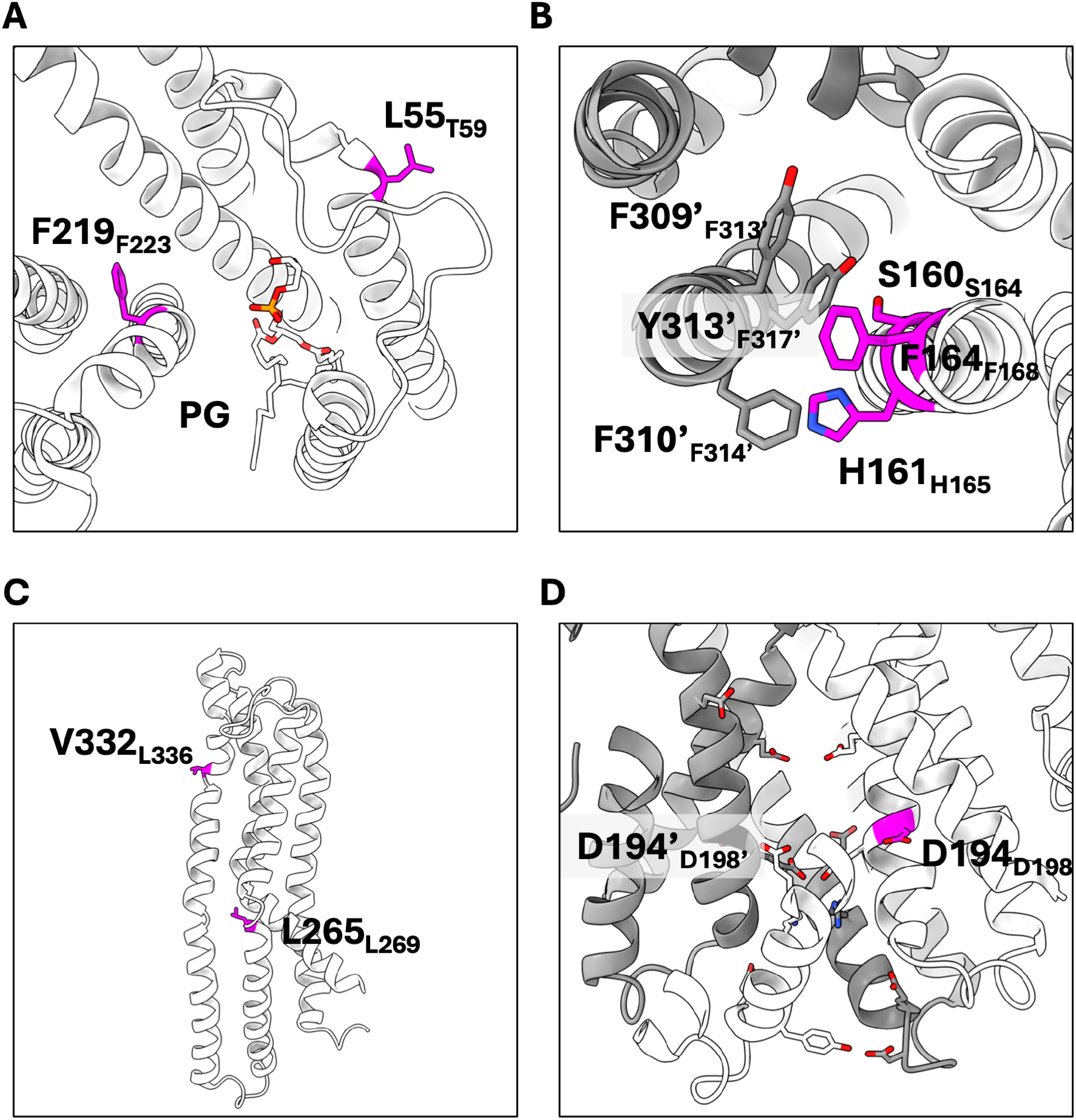
Structure-based analysis of the missense mutations. (**A, B**) Mutants that result in null phenotype and no protein are likely due to the destabilization of the protein. (**A)** The T59C/L55 and F223G/F219 mutants are close to the TM0 domain that may interfere with correct folding. (**B)**. Mutations close to the aromatic rich pocket, Y313’/Y309’, F314’/F310’, and Y309’/Y313’, can disrupt the dimer interface. (**C**,**D**) Mutations that show null phenotype but make protein are likely interfering with conformational changes or proper folding. (**C**) Mutations at the kinks of TM4 and TM5 result in misfolded protein according to the SCAM data as they are important for conformational changes and structural integrity. (**D**) D198G/D194, which is found close to the glutamate ladder and cytoplasmic gate, is being shown to be essential for transport activity and it is likely that the proton cannot be translocated thus rendering the transporter inactive.

### Missense mutations that result in a null phenotype but production of a stable protein

Of the 14 mutants with a null (Fix^-^ NCR247^-^ Bac7^-^) phenotype, 5 make normal or substantial amounts of protein (BacA_*Sm*_ mutant/SbmA_*Ec*_: N113C/N109, D198G/D194, L269C/L265, Y320C/Y316, and L336C/V332). Interestingly, three of these mutants (L269C_Sm_/L265_*Ec*_, Y320C_Sm_/Y316_*Ec*_, and L336C_Sm_/V332_*Ec*_) affect amino acids conserved between BacA and SbmA located in distinctive positions that likely interfere with either conformational changes or correct folding (Fig. 2C). The SCAM data for both the L269C/L265 and L336C/V332 suggests misfolded protein. L269C/L265 is found in the middle of TM4 that displays a kinked conformation. TM4 undergoes conformational changes from outward-to inward-open states and the introduction of a cysteine may be interfering with correct folding. Similarly, the L336C/V332 mutant is located at the top of TM5 where a small helix kink exists. The null phenotype in the D198G/D194 mutant is consistent with our previous work where D194A resulted in reduced to no transport of Microcin B17 and Microcin J25, respectively. D198G/D194 is located at the interface of the glutamate ladder and the cytoplasmic gate (Fig. 2D), and we proposed that the proton might bind to D194 during the transport cycle.

### Missense mutations that result in the production of a stable protein but differentially affect symbiotic nitrogen fixation and transport of the NCR247 and Bac7 peptides

In addition to the 5 *bacA*_*Sm*_ missense mutations discussed above that result in production of a stable protein and cause a null phenotype, 6 missense mutations caused “*split phenotypes”* that strongly differentially affect symbiotic nitrogen fixation and transport of the NCR247 and Bac7 peptides (Fig. 1). These partial loss of function mutants fell into the three different classes: i) **Fix**^**-**^ **NCR247**^**-**^ **Bac7**^**+**^ (N335C/N331). This phenotype could result from a defect in importing NCR247 and possibly some other symbiotically essential NCR peptide(s). while retaining an ability to import the symbiotically irrelevant Bac7 peptide. ii) **Fix**^**-**^ **NCR247**^**+**^ **Bac7**^**+**^ (Y166C/Y162, Q332C/Q328, R389G/Q385, and the central gate residue Y372C/Y368). This phenotype could result from a defect in importing at least one symbiotically essential NCR peptide other than NCR247 while retaining an ability to import the symbiotically irrelevant Bac7 peptide. iii) A **Fix**^**-**^ **NCR247**^**+**^ **Bac7**^**-**^ (F363G/F359), which could result from a defect in importing at least one symbiotically essential NCR peptide(s) other than NCR247 and a defect in importing the symbiotically irrelevant Bac7 peptide. An additional 7 missense mutations resulted in milder split phenotypes (Q193G/Q189, T199C/T195, V206C/L202, G241C/G237, G338C/G334, G349C/G345, and Q365C/Q361).

The structural locations of the 13 missense mutations that caused strong and mild split phenotypes proved to be informative. For 5 of these (Q332C, N335C, F363G, Q365C, and Y372C), the residues that are altered are amino acids that line the cavity in its outward-open conformation. Y372/Y368, the central gate, is at the bottom of the cavity (Fig. 3A-C). In addition, the Fix^-^ NCR247^-^ Bac7^-^ null missense mutant N113C also affects a residue lining the cavity (Fig. 3A-C). Interestingly, except for the Y372 central gate amino acid and N113, these residues line the inside of the cavity in the inward-open conformation as well (Fig. 3D-F). The observation that the two N-terminal residues of Bac7 are important for its import by SbmA_Ec_ (39), suggests that particular residues of NCR247 and other NCR peptides might similarly be of particular importance for their import by BacA_Sm_ and SbmA_E*c*_. In addition, the G241C mutation (in the periplasmic loop between TM3 and TM4) and the G349C mutation (in the periplasmic loop between TM5 and TM6) result in mild split phenotypes (Fig. 3A-B). Taken together, these observations demonstrate that even a simple missense mutation affecting the peptide binding cavity in the open-outward conformation can affect BacA’s ability to discriminate between different substrates.

**Figure 3.**
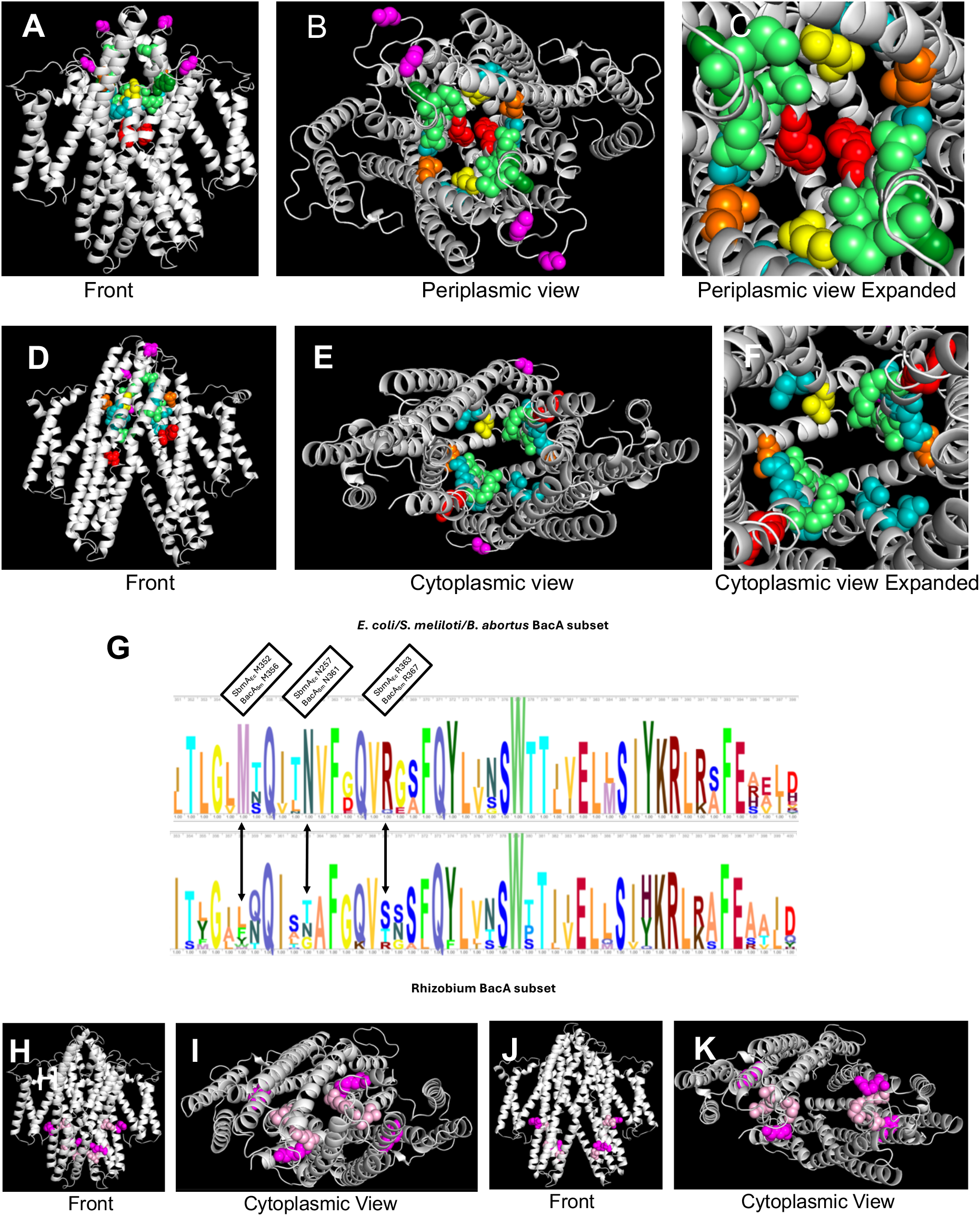
Amino acids lining the BacA cavity in the outward-open conformation that affect the *bacA* phenotype. A-C. Outward-open confromation: Front, Periplasmic, and Expanded Periplasmic views. **D-F**. Inward-open conformation: Front, Cytoplasmic, and Expanded Cytoplasmic views. BacA number and mutation followed by corresponding SbmA number. Residues whose mutation results in a split phenotype: Fix^-^ NCR247^+^ Bac7^+^ (Q332C_*Sm*_/Q328_*Ec*_ and Q365C_*Sm*_/Q361_*Ec*_ teal; Y372C_*Sm*_/Y368_*Ec*_ central gate red); Fix^-^ NCR247^-^ Bac7^+^ (N335C_*Sm*_/N331_*Ec*_ yellow); Fix^-^ NCR247^+^ Bac7^-^ (F363G_*Sm*_/F35 _*Ec*_ forest green). Mutation of another residue lining the cavity, N113C_*Sm*_/(N109_*Ec*_ (orange), results in a null phenotype. In addition, the G241C_*Sm*_/G23 _*Ec*_ mutation (in the loop between TM3 and TM4) and the G349C_*Sm*_/G345_*Ec*_ mutation (in the loop between TM5 and TM6) (magenta) result in mild split phenotypes. Residues M356_*Sm*_/M352_*Ec*_, N361_*Sm*_/N357_*Ec*_, and R367_*Sm*_/R363_*Ec*_ (lime) differ between the BacA_Sm_, SbmA_Ec_, BacA_Ba_ group and the *Rhizobium* BacA group. **G**. A comparison of portions of the amino acid sequence logos of the two BacA subsets developed by diCenzo *et al*. (28) showing that three amino acids (M356/M352, N361/N357, R367/R363 that line the outward-open peptide-binding cavity are not conserved between the BacA_Sm_, SbmA_Ec_, BacA_Ba_ group and the *Rhizobium* BacA group. **H-K**. Amino acids located in the cytoplasmic part of BacA whose mutation results in a strong split phenotype (Y166C_*Sm*_/Y162_*Ec*_ and R389G_*Sm*_ C/385_*Ec*_ magenta) or a mild split phenotype (Q193_*Sm*_G/189G_*Ec*_,T199C_*Sm*_ /T195_*Ec*_, and V206C_*Sm*_ /L202_*Ec*_ pink) **H-I**. Outward-open confromation: Front, Cytoplasmic views. **J-K**. Inward-open conformation: Front, Cytopolasmic views.

A comparison of sequence differences between the BacA_Sm_/SbmA_Ec_/BacA_Ba_ group (restore nitrogen fixation to an *S. meliloti ΔbacA* mutant) and the *Rhizobium* BacA group (do not restore nitrogen fixation to an *S. meliloti ΔbacA* mutant) revealed three residues that stand out as being different between the two classes. These amino acids (BacA_*Sm*_ /SbmA_*Ec*_: M356/M352, N361/N357, and R367/R363) (Fig. 3G) also line the cavity in the outward-open conformation. In addition, N335, whose mutation results in a split phenotype, often differs between the two groups. Fig. 3A-F shows the location of all these residues in the cavity in outward- and inward-open conformations.

Despite the striking number of missense mutations causing a split phenotype affecting the surface of the cavity in the outward-open conformation, we note that amino acid alterations elsewhere in the protein can also result in split phenotypes. Y166C/Y162 (Fix^-^ NCR247^+^ Bac7^+^) and R389G/385 (Fix^-^ NCR247^+^ Bac7^+^) gave very strong split phenotypes, while Q193/Q189, T199/T195, V206/L202 resulted in mild split phenotypes. Both Y166/Y162 and Q193/Q189 line the end of the inward-open cavity and could potentially make direct contact with transported peptide.T199/T195, V206/L202, and make interhelical contacts and could influence the specificity of BacA by distorting the shape of the cavity during peptide import (Fig. 3H-K).

## Discussion

Our detailed genetic, biochemical, structural, and physiological analyses have revealed that BacA_Sm_ and SbmA homodimers function as finely tuned import machines, whose structures can be relatively easily disrupted by single amino acid changes that prevent a stable protein from being made or result in an intact but non-functional protein. Our efforts to relate the location of the missense mutations within the BacA structure of the BacA homodimer to our knowledge of BacA physiology and its evolution suggested a plausible explanation for one of the most striking but perplexing results concerning BacA’s role in symbiosis - *bacA* orthologs from two mammalian pathogens (SbmA_Ec_ and BacA_Ba_) can fully restore the ability of an *S. meliloti ΔbacA* mutant to establish a normal nitrogen-fixing symbiosis with its *Medicago hosts* M. *truncatula* and *M. sativa* (26, 27), but *bacA* orthologs from rhizobial strains (e.g. *S. fredii* NGR234, *R. leguminosarum* bv. *viciae* and *sativum*) that interact with different legume hosts cannot (28, 29). These rhizobial *bacA* orthologs do, however, allow a *S. meliloti ΔbacA* mutant to proceed to a somewhat more developmentally advanced stage than is observed in *ΔbacA* nodules although many of the cells eventually die.

Our analyses suggest that single amino acid changes, particularly in the proposed peptide-binding cavity in the outward-facing conformation, can play roles in the rapid co-evolution of nitrogen-fixing rhizobia with their legume partners. Our previous phylogenetic analyses (36) indicated that SbmA/BacA likely evolved from the transmembrane domain (TMD) of the ABC transporter YddA, which in some organisms is expressed from a single gene but in others from two separate genes encoding TMD and ABC subunits. The proposed evolutionary mechanism is that a gene encoding only the TMD domain was horizontally transferred to a new organism, where it subsequently evolved a glutamate ladder and other structural adaptions that allowed it to be powered by the membrane potential instead of ATP hydrolysis. On the basis of their analyses, Smith *et al*. (40) suggest that the SbmA/BacA family originated in a bacterium in the order Rhizobiales over 500 million years ago prior to spreading to other Proteobacteria by horizontal gene transfer. Since legumes did not evolve until about 60 million years ago (41), the initial spread of SbmA/BacA family members may have been driven in part by their abilities to protect against membrane-damaging antimicrobial peptides, a trait beneficial to animal and plant pathogens, and that they were only subsequently co-opted to support legume symbiosis in rhizobia.

Since some legumes began to express NCR peptides in their nodules to increase the nitrogen-fixing efficiency of the symbiosis, both these NCR peptide families and the NCR importer BacA in their symbiotic rhizobial partner have undergone rapid co-evolution (42). Our analyses suggest that one important evolutionary mechanism that allows a BacA to retain its relatively promiscuous import ability yet finely adapt to the cocktail of NCR peptides made by its particular legume host (4) is to make amino acid changes in outward-facing peptide binding cavity of BacA. Since NCR peptides appear to have rapidly evolved and diversified (43, 44), there has been a corresponding rapid evolution in the substrate specificity of the BacA proteins (28), which is likely achieved at least in part by subtle modifications to the peptide-binding cavity. Due to the substantial number of highly cationic NCR peptides encoded by its hosts, *M. truncatula and M. sativa, S. meliloti’s* BacA appears particularly adept at importing such cationic peptides (29). This line of reasoning can explain the incomplete complementation following expression of BacAs in rhizobia whose legume hosts have a lower proportion of highly cationic NCR peptides. Moreover, *S. meliloti* BacA has undergone rapid, convergent evolution to that of the BacA/SbmA proteins of the pathogenic genera *Brucella, Escherichia, and Klebsiella* (28), perhaps driven by the exposure of those pathogens to highly cationic peptides. Thus, this line of reasoning can also explain the otherwise paradoxical observation that BacA/SbmA orthologs from the pathogens *E. coli* and *B. abortus* can make an *S. meliloti bacA* mutant fully symbiotically proficient, whereas BacA orthologs from numerous other rhizobia cannot. NCR peptides are expressed in consecutive waves during infection of wild-type nodules and can be separated into early and late stage NCR peptides (4). it is possible that the ability of these rhizobial *bacA* orthologs to nevertheless allow a *S. meliloti ΔbacA* mutant to proceed to a somewhat more developmentally advanced stage than is observed in *ΔbacA* nodules could be due them being able to import NCR peptides expressed early in nodule development, but not certain cationic NCR peptides such as NCR247 expressed during late nodule development.

## Materials and Methods

BacA protein production, BacA antibody generation, the protocols for the Substituted Cysteine Accessibility Method (SCAM), and Protein Analysis of BacA are described in the SI Materials and Methods.

### Bacterial strains

The bacterial strains used in this study are described in Table S1 and their growth conditions in SI Materials and Methods

### BacA cloning and site directed mutagenesis

To create defined site directed mutations (SDM), the *S. meliloti bacA* gene was cloned into the EcoRI and BamHI sites of pUC19. On this construct, Site directed mutagenesis of this cloned gene was performed using a QuikChange II site directed mutagenesis kit (Agilent), according to the manufacturer’s instructions. Primers used for SDM creation of the defined SDMs are shown in Table S2.

### BacA complementation constructs

All the *bacA* complementation constructs that were generated for this study are shown in Table S3. Their construction is described in SI Materials and Methods.

### Assays to determine phenotype of *bacA* mutants

*Medicago sativa* plant symbiosis experiments were conducted exactly as previously described (45). NCR247 sensitivity survival assays were performed in MOPS buffer supplemented with casamino acids as described previously using the defined concentrations of NCR247 (21). In some cases, the incubation time was extended to 10 hours to observe better separation between wild type/complemented and Δ*bacA* mutant bacteria. Bac7(1-35) sensitivity survival assays were performed as described previously (34).

## Acknowledgments

This work was supported by NIH grants (R01 GM031030) to G.C.W., Imperial - BBSRC International Partnership Fund to K.B. and G.C.W. S.S was funded by the Stowers Institute for Medical Research. We thank Barbara Imperiali for her assistance with the SCAM method. G.C.W. is an American Cancer Society Professor

