## Supporting Information for "The BacA(SbmA) Importer of Symbiotically Important Legume Nodule Cysteine-Rich Peptides: Insights into Protein Architecture, Function, and Evolutionary Implications"

**This PDF file includes:**

Materials and Methods
Figures S1-S4
Table S1-4
References

### Materials and Methods

**Bacterial strains.** The bacterial strains used in this study are described in Table S1. *S. meliloti* strains were grown in Luria-Bertani broth (LB) alone or in LB supplemented with 2.5 mM $CaCl_2$ and 2.5 mM $MgSO_4$ (LB-MC) (1) for 48 h. When required the growth medium was supplemented with 500 ug/ml of streptomycin and 10 ug/ml of Tetracycline when necessary. *E. coli* cells were grown in LB supplemented with 10 ug/ml of Tetracycline, when required.

#### BacA cloning and site directed mutagenesis

To create defined site directed mutations (SDM), the *SmbacA* gene was cloned into the EcoRI and BamHI sites of pUC19. On this construct, site directed mutagenesis was done using a QuikChange II site directed mutagenesis kit (Agilent), according to the manufacturers' instructions. Primers used for SDM creation of the defined SDMs are shown in Table S2.

#### BacA complementation constructs

For complementation for the loss of *bacA* in *S. meliloti,* the mutated *S. meliloti bacA* genes were PCR amplified from the mutated pUC19-SmBacA plasmids, using the primers: *Smeliloti_bacA*_NsiI_RBS_NcoI_F and *Smeliloti_bacA*_XbaI_R followed by column purification of the PCR product. The *bacA* genes were then cloned into the PstI and XbaI sites of pRF771, which placed them under the control of a constitutive *trp* promoter. All cloning with pRF771 was perfomed using plasmids, isolated from a *dam-/dcm- E. coli* strain, GM272. The resulting constructs were transformed into E. coli Dh5α and sequence verified with primers SmbacA_seq_F and SmbacA_seq_R by Sanger sequencing. Then the newly generated pBacASDM constructs were transformed into *Sm*1021 Δ*bacA* by triparental mating as previously described (2), All the *bacA* complementation constructs that were generated for this study are shown in Table S3.

#### BacA protein production

BacA for polyclonal antibodies was prepared as previously described without any modifications ((3, 4). In brief, BacA was overexpressed in Lemo21 (DE3) cells and purified in 20 mM tris (pH 8.0), 150 mM NaCl, and 0.03% n-Dodecyl β-D-maltoside (DDM).

#### BacA antibody generation

Antibodies against BacA were raised using two rabbits. We used a 118-day protocol with Covance Research Products, an AALAC certified company. Our CAC protocol numbers is 0413-029-16. All personnel handling animals were CAC certified.

#### Substituted Cysteine Accessibility Method (SCAM)

SCAM assays were performed as previously described with modifications (5). Sm1021 cultures were grown in LBMC with selection antibiotics (Strep and Tet). Cultures were then diluted into fresh LB (with the same selection antibiotic as above) to a final $OD_{600}$ of 0.2 and then washed once with PBS (137 mM NaCl, 2.7 mM KCl, 10 mM $Na_2HPO_4$, 1.8 mM $KH_2PO_4$, pH 7.4), adjusted to a final $OD_{600}$ of 1, and divided into four 50 µL aliquots. Aliquots received 5 µl of 55 mM N-ethylmaleimide (NEM, Alfa-Aesar #40526) or sodium(2-sulfonatoethyl)methanethiosulfonate (MTSES, Cayman Chemical #16529) as indicated for a final concentration of 5 mM and were incubated at room temperature in the dark for 1h for labeling to proceed. Cells were harvested by centrifugation at 16,000x*g* for 3 min and washed once with 100 µl of PBS. Pellets were re-suspended in 22.5 µl of lysis buffer (50 mM HEPES, 5% SDS, pH 7.5) and 7.5 µl of 25 mM PEG-mal (Sigma #63187) in DMSO. Samples were mixed well and incubated at room temperature in the dark for 1-1.5 h. Reactions were quenched by the addition of 30 µl of 2x loading buffer (100 mM Tris pH 6.8, 10% glycerol, 3% SDS, 6 M urea, 1% β-mercaptoethanol, 0.1% bromophenol blue) and frozen at -20 ºC until needed.

#### Protein analysis BacA

Proteins were separated by SDS-PAGE (3 µl per lane) on 12% acrylamide gels alongside BioRad Precision Plus standards and transferred to nitrocellulose membranes for Western blotting with the BacA specific antibodies obtained for this study. Blots were blocked with 5% Milk in TBST (10 mM Tris, 150 mM NaCl, 0.05% Tween-20, pH 7.5) for 1 hour. The primary antibody, rabbit anti BacA was applied in a

1:10,000 dilution for 1 h. The secondary antibody, goat anti-rabbit-HRP was applied in a 1:25,000 dilution for 1 h.

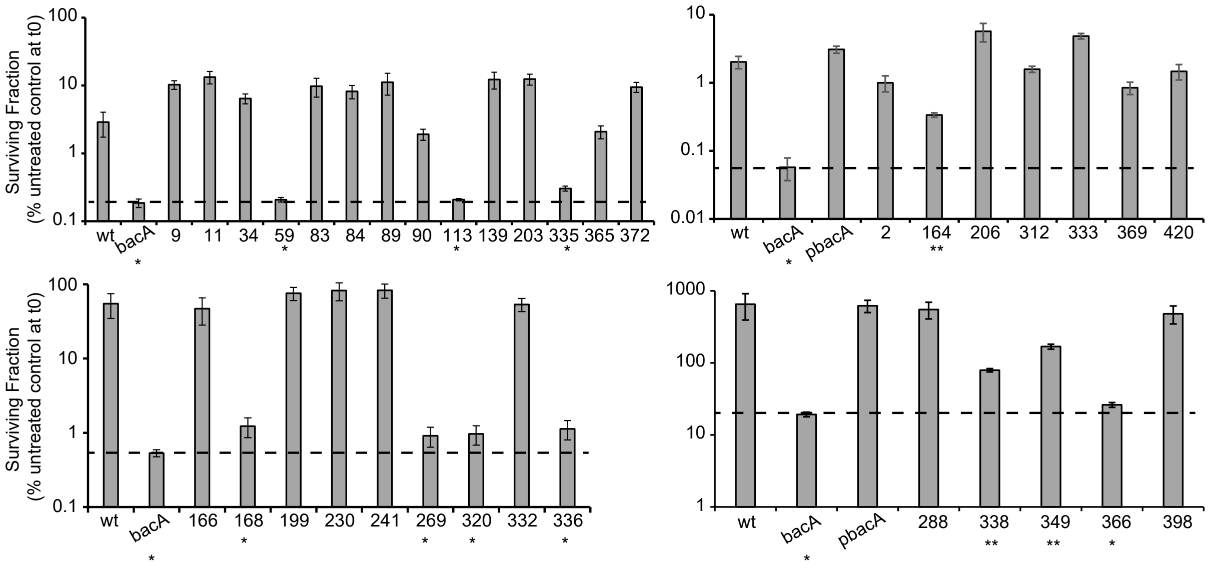


**Fig. S1. Sensitivity of *bacA* mutants to killing by NCR247**.*S. meliloti Sm*1021 cells were treated with 20 µM NCR247 for 180 min in MOPS-GS buffer supplemented with 1% casamino acids. Bars show the relative survival of Sm1021 bacA mutant strains carrying either the pRF771 control plasmid or pRF771with the defined BacA variants. All error bars represent mean ± SD (*n* = 3) based on three technical replicates representing trends observed in at least two independent experiments. * - BacA variant does not protect against NCR247, ** - BacA variant does partially protect against NCR247.

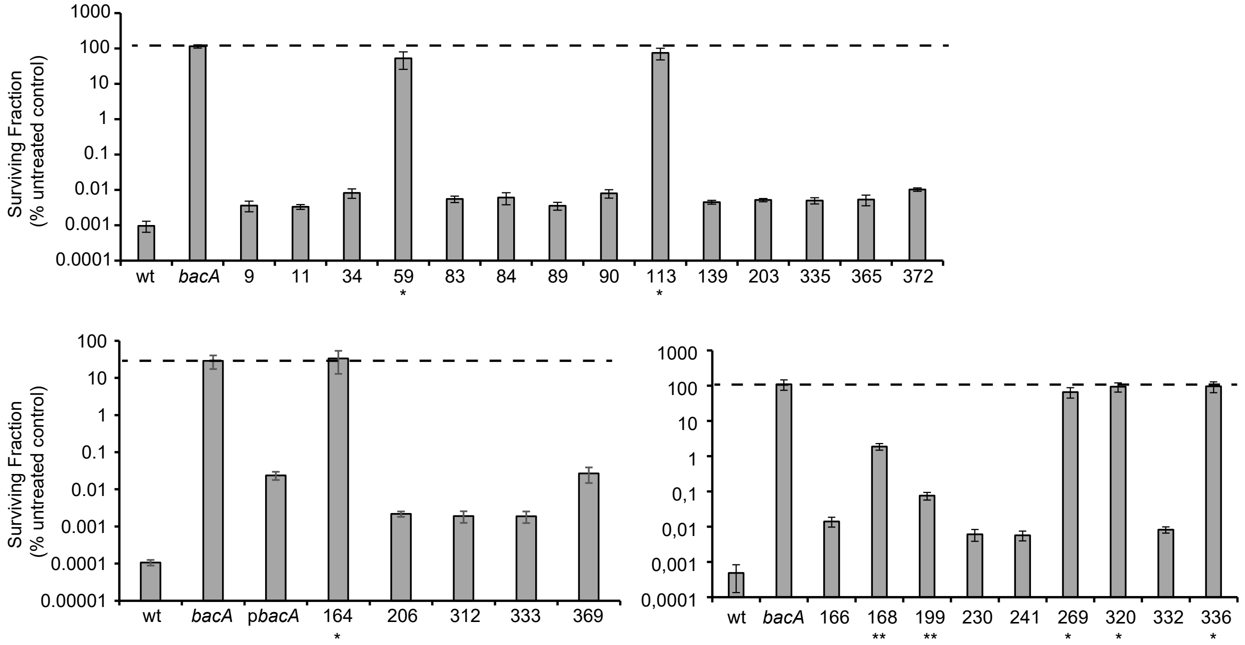


**Fig. S2. Sensitivity of *bacA* mutants to killing by Bac7 (1-35).** Cultures of *S. meliloti Sm*1021 were exposed to 1 µM Bac7(1-35), and cell viability was determined in LB relative to an untreated control after 1 hour. Bars show the relative survival of *Sm*1021 bacA mutant strains carrying either the pRF771 control plasmid or pRF771 with the defined BacA variants*.* All error bars represent mean ± SD (*n* = 3) based on three technical replicates representing trends observed in at least two independent experiments. * - BacA variant does not protect against NCR247, ** - BacA variant does partially protect against NCR247.

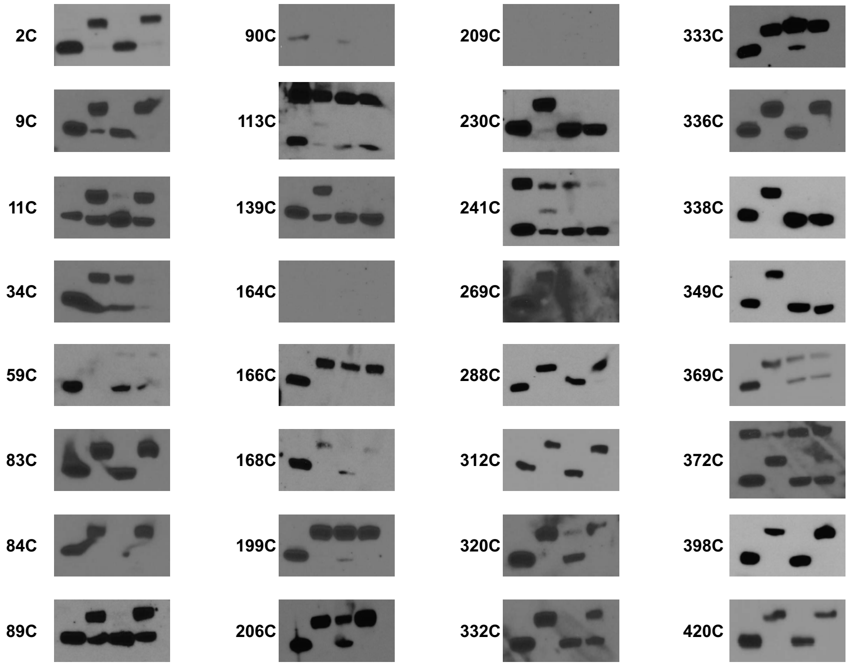


**Fig. S3. Behavior of the monocysteine *bacA* mutants that made protein in the SCAM analysis for determining the topology of membrane proteins *in vivo*.** Aliquots received either the membrane permeable alkylating agent N-ethylmaleimide (NEM) to react with any thiol from a cysteine residue exposed at either the cytoplasmic or the periplasmic interface of BacA or the membrane impermeable agent sodium(2-sulfonatoethyl)methanethiosulfonate (MTSES) to react with any thiol group of a cysteine exposed at the periplasmic interface of BacA. After lysis the high molecular weight compound PEG maleimide (PEG-mal) was added to react with any unreacted thiol groups. Due to the high molecular weight of PEG-mal a shift of molecular weight of BacA can be detected. For each gel depiction the samples were loaded in the following order: 1. Untreated control, 2. PEG-mal control, 3. Membrane permeable NEM, 4. Membrane impermeable MTSES.

```
     1        10        20        30        40        50        60
BacA MFQSFFPKPKLFFISSAVWSLLAVLAWYAGGRDIGAYLGLPPLPPGQEPVIGVSVFWSTP
SbmA MFKSFFPKPGTFFLSAFVWALIAVIFWQAGGGDWVARIT...GASGQI.PISAARFWSLD

              70        80        90       100       110       120
BacA FLWFYIYYAVVAGLFAAFWFAYSPHRWQYWSVLGTALIIFNTYFSVQVSVAINAWYGPFY
SbmA FLIFYAYYIVCVGLFALFWFIYSPHRWQYWSILGTALIIFVTWFLVEVGVAVNAWYAPFY

             130       140       150       160       170       180
BacA DLIQQALARTAPVTAGQLYSGMIGFSGIAFVAVTVGVLNLFFVSHYIFRWRTAMNEFYVA
SbmA DLIQTALSSPHKVTIEQFYREVGVFLGIALIAVVISVLNNFFVSHYVFRWRTAMNEYYMA

             190       200       210       220       230       240
BacA HWPRLRHVEGASQRVQEDTMRFSSTVERLGVGLVSSIMTLIAFLPVLFKFSEQVNVLPIV
SbmA NWQQLRHIEGAAQRVQEDTMRFASTLENMGVSFINAIMTLIAFLPVLVTLSAHVPELPII

             250       260       270       280       290       300
BacA GEIPHALVWAAVFWSVFGTVFLAAVGIKLPGLEFRNQRVEAAYRKELVYGEDHEDRADPI
SbmA GHIPYGLVIAAIVWSLMGTGLLAVVGIKLPGLEFKNQRVEAAYRKELVYGEDDATRATPP

             310       320       330       340       350       360
BacA TLAQLFDNVRRNYFRLYFHYMYFNIARIFYLQADNLFGTFVLVPAIVAGKLTLGVMNQVL
SbmA TVRELFSAVRKNYFRLYFHYMYFNIARILYLQVDNVFGLFLLFPSIVAGTITLGLMTQIT

             370       380       390       400       410       420
BacA NVFGQVRESFQYLVNSWTTIVELLSIYKRLKAFESVLVDEPLPEIDRQFIDAGGKEELAL
SbmA NVFGQVRGAFQYLINSWTTLVELMSIYKRLRSFEHELDGDKIQEVTHTLS..........
```

**Fig. S4. Sequence alignment of SbmA (*E. coli*, Uniprot P0AFY6) and BacA (*S. meliloti,* Uniprot Q08120).** Identical residues as indicated with red fill, while conservative substitutions are indicated in red letters.

**Table S1. Bacterial strains used in this study**

***S. meliloti***

| ***Bacterial strain*** | ***Relevant characteristics*** | **Source** |
|---|---|---|
| *Sm*1021 wild type | Smr derivative of SU47 | (Meade *et al.*, 1982) |
| *Sm*1021 Δ*bacA* | Sm1021, *bacA654*::SpcR | (Ferguson *et al.*, 2002) |

***Escherichia coli***

| **Bacterial Strain** | **Relevant characteristics** | **Source** |
|---|---|---|
| DH5α | *supE44 lacU169 (80lacZ M15) hsdR17 recA1 endA1 gyrA96 thi-1 relA1 luxS* | Bethesda Research Laboratories |
| GM272 | *hsd*S2I *dam*-3 *dcm*-6 *E. coli* | Genetic Stock Centre |
| MT616 | MM294A *recA56* (pRK600) Cmr | (Finan *et al.*, 1986) |

**Table S2. Primer sequences used in this study BacA cloning and site-directed mutagenesis**

.

| Primer name | sequence |
|---|---|
| BacASDMV288C_F | GCATACCGGAAGGAACTGTGCTATGGCGAGGACCACGAA |
| BacASDMV288C_R | TTCGTGGTCCTCGCCATAGCACAGTTCCTTCCGGTATGC |
| BacASDMG338C_F | CAGGCCGACAATCTCTTCTGCACTTTCGTGCTGGTCCCG |
| BacASDMG338C_R | CGGGACCAGCACGAAAGTGCAGAAGAGATTGTCGGCCTG |
| BacASDMG349C_F | GTCCCGGCAATCGTGGCGTGTAAGCTCACACTCGGCGTC |
| BacASDMG349C_R | GACGCCGAGTGTGAGCTTACACGCCACGATTGCCGGGAC |
| BacASDMV366C_F | CTGAATGTGTTCGGGCAGTGCCGGGAGTCCTTCCAGTAT |
| BacASDMV366C_R | ATACTGGAAGGACTCCCGGCACTGCCCGAACACATTCAG |
| BacASDMV398C_F | GCCTTTGAGTCGGTATTGTGCGACGAGCCGCTGCCGGAA |
| BacASDMV398C_R | TTCCGGCAGCGGCTCGTCGCACAATACCGACTCAAAGGC |
| BacASDML420C_F | GGTAAGGAAGAGCTGGCTTGCTAACGGCGGGCATCAGAA |
| BacASDML420C_R | TTCTGATGCCCGCCGTTAGCAAGCCAGCTCTTCCTTACC |
| BacASDML001C_F | AAACGACGGCCAGTGAATTCTGCTTCCAATCCTTCTTCC |
| BacASDML001C_R | GGAAGAAGGATTGGAAGCAGAATTCACTGGCCGTCGTTT |
| BacASDMS164C_F | GCTCAACCTCTTTTTCGTCTGCCATTACATCTTTCGCTG |
| BacASDMS164C_R | CAGCGAAAGATGTAATGGCAGACGAAAAAGAGGTTGAGC |
| BacASDMV206C_F | CGTTTCTCCAGCACATGCGAGAGGCTGGGTGTTGGTCT |
| BacASDMV206C_R | AGACCAACACCCAGCCTCTCGCATGTGCTGGAGAAACG |
| BacASDMN312C_F | TTCGACAATGTCAGGCGCTGCTATTTTCGCCTGTATTTCC |
| BacASDMN312C_R | GGAAATACAGGCGAAAATAGCAGCGCCTGACATTGTCGAA |
| BacASDMA333C_F | CCGCATTTTCTACCTGCAGTGCGACAATCTCTTCGGCACT |
| BacASDMA333C_R | AGTGCCGAAGAGATTGTCGCACTGCAGGTAGAAAATGCGG |
| BacASDMS369C_F | TGTTCGGGCAGGTGCGGGAGTGCTTCCAGTATCTCGTCA |
| BacASDMS369C_R | TGACGAGATACTGGAAGCACTCCCGCACCTGCCCGAACA |
| BacASDML420C_F | GGTAAGGAAGAGCTGGCTTGCTAAGGATCCTCTAGAGTCG |
| BacASDML420C_R | CGACTCTAGAGGATCCTTAGCAAGCCAGCTCTTCCTTACC |
| BacASDMF002C_F | GACGGCCAGTGAATTCTTGTGCCAATCCTTCTTCCCCAAG |
| BacASDMF002C_R | CTTGGGGAAGAAGGATTGGCACAAGAATTCACTGGCCGTC |
| BacASDMP009C_F | CAATCCTTCTTCCCCAAGTGCAAGCTGTTTTTCATATCTTCCGCC |
| BacASDMP009C_R | GGCGGAAGATATGAAAAACAGCTTGCACTTGGGGAAGAAGGATTG |
| BacASDML011C_F | CCTTCTTCCCCAAGCCGAAGTGCTTTTTCATATCTTCCGCCG |
| BacASDML011C_R | CGGCGGAAGATATGAAAAAGCACTTCGGCTTGGGGAAGAAGG |
| BacASDMI034C_F | TATGCCGGGGGGCGGGACTGCGGCGCCTACCT |
| BacASDMI034C_R | AGGTAGGCGCCGCAGTCCCGCCCCCCGGCATA |
| BacASDMT59C_F | GCGTGTCCGTTTTCTGGTCCTGCCCTTTTCTTTGGTTTTAC |
| BacASDMT59C_R | GTAAAACCAAAGAAAAGGGCAGGACCAGAAAACGGACACGC |
| BacASDMS083C_F | TTCTGGTTCGCCTACTGCCCGCATCGGTGGCAGTATT |
| BacASDMS083C_R | AATACTGCCACCGATGCGGGCAGTAGGCGAACCAGAA |
| BacASDMP084C_F | TTCTGGTTCGCCTACAGCTGCCATCGGTGGCAGTATTGGTCG |
| BacASDMP084C_R | CGACCAATACTGCCACCGATGGCAGCTGTAGGCGAACCAGAA |

| | |
|---|---|
| BacASDMY089C_F | CATCGGTGGCAGTGCTGGTCGGTACTCGGCA |
| BacASDMY089C_R | TGCCGAGTACCGACCAGCACTGCCACCGATG |
| BacASDMW090C_F | ATCGGTGGCAGTATTGCTCGGTACTCGGCA |
| BacASDMW090C_R | TGCCGAGTACCGAGCAATACTGCCACCGAT |
| BacASDMN113C_F | TCAGCGTCGCGATCTGCGCGTGGTACGGCCCTTT |
| BacASDMN113C_R | AAAGGGCCGTACCACGCGCAGATCGCGACGCTGA |
| BacASDMY139C_F | ACAGCAGGGCAGCTCTGCTCGGGAATGATCGGATT |
| BacASDMY139C_R | AATCCGATCATTCCCGAGCAGAGCTGCCCTGCTGT |
| BacASDMY166C_F | CTCTTTTTCGTCAGCCATTGCATCTTTCGCTGGCGCAC |
| BacASDMY166C_R | GTGCGCCAGCGAAAGATGCAATGGCTGACGAAAAAGAG |
| BacASDMF168C_F | TTCGTCAGCCATTACATCTGCCGCTGGCGCACGG |
| BacASDMF168C_R | CCGTGCGCCAGCGGCAGATGTAATGGCTGACGAA |
| BacASDMF230C_F | TCTCCCGGTACTCTTCAAGTGCTCCGAACAGGTGAAC |
| BacASDMF230C_R | GTTCACCTGTTCGGAGCACTTGAAGAGTACCGGGAGA |
| BacASDMG242C_F | AACGTCCTGCCCATTGTCTGCGAGATACCGCA |
| BacASDMG242C_R | TGCGGTATCTCGCAGACAATGGGCAGGACGTT |
| BacASDML269C_F | CCGCAGTCGGCATCAAGTGCCCGGGACTGGAATTTC |
| BacASDML269C_R | GAAATTCCAGTCCCGGGCACTTGATGCCGACTGCGG |
| BacASDMQ332C_F | ATTGCCCGCATTTTCTACCTGTGCGCCGACAATCTCTTCGGCA |
| BacASDMQ332C_R | TGCCGAAGAGATTGTCGGCGCACAGGTAGAAAATGCGGGCAAT |
| BacASDMQ365C_F | TCCTGAATGTGTTCGGGTGCGTGCGGGAGTCCTTCCA |
| BacASDMQ365C_R | TGGAAGGACTCCCGCACGCACCCGAACACATTCAGGA |
| BacASDMY372C_F | TGCGGGAGTCCTTCCAGTGCCTCGTCAATTCCTG |
| BacASDMY372C_R | CAGGAATTGACGAGGCACTGGAAGGACTCCCGCA |
| BacASDMY320C_F | GCCTGTATTTCCATTACATGTGCTTCAATATTGCCCGCAT |
| BacASDMY320C_R | ATGCGGGCAATATTGAAGCACATGTAATGGAAATACAGGC |
| BacASDMS203C_F | AGGACACGATGCGTTTCTGCAGCACAGTCGAGAGG |
| BacASDMS203C_R | CCTCTCGACTGTGCTGCAGAAACGCATCGTGTCCT |
| BacASDMT199C_F | CAGCGTGTGCAAGAGGACTGCATGCGTTTCTCCAGCACAGT |
| BacASDMT199C_R | ACTGTGCTGGAGAAACGCATGCAGTCCTCTTGCACACGCTG |
| BacASDMN335C_F | TCTACCTGCAGGCCGACTGCCTCTTCGGCACTTTCGTGCT |
| BacASDMN335C_R | AGCACGAAAGTGCCGAAGAGGCAGTCGGCCTGCAGGTAGA |
| BacASDML336C_F | ACCTGCAGGCCGACAATTGCTTCGGCACTTTCGTGCTG |
| BacASDML336C_R | CAGCACGAAAGTGCCGAAGCAATTGTCGGCCTGCAGGT |
| SmbacA_seq_F | GATGAACGAATTTTATGTCGCGCATTGGC |
| SmbacA_seq_R | GGGCAATATTGAAGTACATGTAATGGAAATACAGGC |
| SmbacA_NsiI_RBS_F | CTAGAATGCATGAAACGAGAGTGCCGTCCCCCTTGTTCCAATCCTTCTT CCCCAAGC |
| SmbacA_His_XbaI_R | TTATCGTCTAGATTACAGAGCCAGCTCTTCCTTACCG |

**Table S3. BacA complementation constructs used in this study**.

| Plasmid name | Relevant characteristics | source |
|---|---|---|
| pRF771 | RK2 derivative P*trp* expression vector Tetr | (6) |
| pUC19 | High copy bacterial cloning vector | (7) |
| pUC19-*SmbacA* | pUC19 carrying the entire *S. meliloti* bacA gene | (8) |
| p*SmbacA* | pRF771 carrying the *S. meliloti bacA* gene under control of a constitutive P*trp* promoter, Tetr | This study |
| **Cysteine mutant** | | |
| pBacASDMV288C | p*SmbacA* carrying the V288C mutation | This study |
| pBacASDMG338C | p*SmbacA* carrying the G338C mutation | This study |
| pBacASDMG349C | p*SmbacA* carrying the G349C mutation | This study |
| pBacASDMV366C | p*SmbacA* carrying the V366C mutation | This study |
| pBacASDMV398C | p*SmbacA* carrying the V398C mutation | This study |
| pBacASDML420C | p*SmbacA* carrying the L420C mutation | This study |
| pBacASDML001C | p*SmbacA* carrying the L001C mutation | This study |
| pBacASDMS164C | p*SmbacA* carrying the S164C mutation | This study |
| pBacASDMV206C | p*SmbacA* carrying the V206C mutation | This study |
| pBacASDMN312C | p*SmbacA* carrying the N312C mutation | This study |
| pBacASDMA333C | p*SmbacA* carrying the A333C mutation | This study |
| pBacASDMS369C | p*SmbacA* carrying the S369C mutation | This study |
| pBacASDMF002C | p*SmbacA* carrying the F002C mutation | This study |
| pBacASDMP009C | p*SmbacA* carrying the P009C mutation | This study |
| pBacASDML011C | p*SmbacA* carrying the L011C mutation | This study |
| pBacASDMI034C | p*SmbacA* carrying the I034C mutation | This study |
| pBacASDMT59C | p*SmbacA* carrying the T59C mutation | This study |
| pBacASDMS083C | p*SmbacA* carrying the S083C mutation | This study |
| pBacASDMP084C | p*SmbacA* carrying the P084C mutation | This study |
| pBacASDMY089C | p*SmbacA* carrying the Y089C mutation | This study |
| pBacASDMW090C | p*SmbacA* carrying the W090C mutation | This study |
| pBacASDMN113C | p*SmbacA* carrying the N113C mutation | This study |
| pBacASDMY139C | p*SmbacA* carrying the Y139C mutation | This study |
| pBacASDMY166C | p*SmbacA* carrying the Y166C mutation | This study |
| pBacASDMF168C | p*SmbacA* carrying the F168C mutation | This study |
| pBacASDMF230C | p*SmbacA* carrying the F230C mutation | This study |
| pBacASDMG242C | p*SmbacA* carrying the G242C mutation | This study |
| pBacASDML269C | p*SmbacA* carrying the L269C mutation | This study |
| pBacASDMQ332C | p*SmbacA* carrying the Q332C mutation | This study |
| pBacASDMQ365C | p*SmbacA* carrying the Q365C mutation | This study |
| pBacASDMY372C | p*SmbacA* carrying the Y372C mutation | This study |
| pBacASDMY320C | p*SmbacA* carrying the Y320C mutation | This study |
| pBacASDMS203C | p*SmbacA* carrying the S203C mutation | This study |
| pBacASDMT199C | p*SmbacA* carrying the T199C mutation | This study |
| pBacASDMN335C | p*SmbacA* carrying the N335C mutation | This study |
| pBacASDML336C | p*SmbacA* carrying the L336C mutation | This study |
| **Glycine mutants** | | |
| pRK404 | Broad host range vector; Tetr | (9) |
| pJG51A | pRK404 carrying the *S. meliloti bacA* gene | (1) |
| W57G | pRK404 with bacA(*W57G*) | (10) |

| | | |
|---|---|---|
| S83G | pRK404 with bacA(*S83G*) | (10). |
| W87G | pRK404 with bacA(*W87G*) | (10) |
| Y120G | pRK404 with bacA(*Y120G*) | (10) |
| N159G | pRK404 with bacA(*N159G*) | (10) |
| H165G | pRK404 with bacA(*H165G*) | (10). |
| W182G | pRK404 with bacA(*W182G*) | (10) |
| Q193G | pRK404 with bacA(*Q193G*) | (10) |
| R194G | pRK404 with bacA(*R194G*) | (10) |
| D198G | pRK404 with bacA(*D198G*) | (10). |
| F223G | pRK404 with bacA(*F223G*) | (10) |
| S231G | pRK404 with bacA(*S231G*) | (10). |
| T259G | pRK404 with bacA(*T259G*) | (10). |
| R284G | pRK404 with bacA(*R284G*) | (10). |
| N312G | pRK404 with bacA(*N312G*) | (10). |
| Q332G | pRK404 with bacA(*Q332G*) | (10) |
| K350G | pRK404 with bacA(*K350G*) | (10). |
| F363G | pRK404 with bacA(*F363G*) | (10) |
| R389G | pRK404 with bacA(*R389G*) | (10) |

**Table S4. Assessment of monocysteine *bacA* mutants for their ability to support plant growth when expressed in an *Sm*1021 *ΔbacA* mutant**

| BacA mutant | Plant Phenotype |
|---|---|
| pRF771 (*bacA*$^-$) | - |
| pSmbacA (*bacA*$^+$) | +++ |
| F2C | ++ |
| P9C | +++ |
| L11C | +++ |
| I34C | +++ |
| T59C | - |
| S83C | +++ |
| P84C | +++ |
| Y89C | +++ |
| W90C | +++ |
| N113C | - |
| Y139C | ++ |
| S164C | - |
| Y166C | - |
| F168C | - |
| T199C | +++ |
| S203C | +++ |
| V206C | +++ |
| F230C | ++ |
| G241C | + |
| L269C | - |
| V288C | ++ |
| N312C | +++ |
| Y320C | - |
| Q332C | - |
| A333C | ++ |
| N335C | + |
| L336C | - |
| G338C | - |
| G349C | ++ |
| Q365C | ++ |
| V366C | - |
| S369C | +++ |
| Y372C | + |
| V398C | +++ |
| L420C | +++ |

Four plants per inoculation were assessed for plant growth and nodule formation as compared to an $H_2O$ inoculation without bacteria. Plants inoculated with wild-type *Sm*1021 did show significant growth with green leaves and multiple pink nodules, on average, as compared to plants inoculated with either the

*Sm*1021 *bacA* deletion mutant or $H_2O$ which showed no significant growth with yellow/brownish leaves and no pink root nodules. Plant phenotypes for the BacA monoglycine variants are given in Levier *et al.* (10).

| | |
|---|---|
| - | No significant plant growth compared to an H2O inoculation without bacteria |
| + | Plants are small but show light green leaves |
| ++ | Plants show some growth and a mix of light and dark green leaves |
| +++ | Plants show growth comparable to the *bacA*- strain complemented with wild-type *bacA* |
